# Adolescent food insecurity impairs gut signal sensitivity and cue-induced appetitive behaviours in female rats

**DOI:** 10.64898/2026.04.29.721762

**Authors:** Abbey Livermore, Zhi Yi Ong

## Abstract

Unpredictable and insufficient access to food, known as food insecurity, is associated with the development of obesity. However, causal mechanisms underlying this paradoxical relationship remain poorly understood. Using a rat model of food insecurity, this study investigated whether food insecurity causes dysregulated feeding behaviours, specifically impaired gut signal sensitivity and enhanced cue-driven appetitive responses. Adolescent female rats were assigned to receive either *ad libitum* chow access (Food secure), 90% caloric restriction (Food restricted) or unpredictable quantity and timing of food access (Food insecure), for 4 weeks. After which, rats were returned to an *ad libitum* chow diet for the remainder of the study. To examine gut signal sensitivity, we measured the effects of cholecystokinin (CCK) on 10% sucrose intake. To examine cue-driven feeding behaviours, we used Pavlovian appetitive conditioning and measured appetitive responses towards a food-predictive cue. Results showed that prior food insecure rats were less sensitive to the intake inhibitory effects of CCK and exhibited enhanced cue-induced appetitive behaviours, when compared to food secure and food restricted groups. Anxiety-like behaviours or learning and memory was not different between groups. At the end of the study, adolescent caloric restriction resulted in reduced fat mass, plasma leptin levels and body weight when compared to food secure, but not food insecure rats, suggesting that adolescent food insecurity somewhat overcame these metabolic effects. Taken together, our findings suggest that adolescent food insecurity impaired gut signal sensitivity and heightened food cue sensitivity, which may cause enduring metabolic and behavioural adaptations that promote overeating and weight gain.

## 1. Introduction

Food insecurity is a state of having unpredictable access to food of adequate quality or quantity to support a healthy lifestyle (Food and Agriculture Organization of the United Nations [FAO] et al., 2023). In 2024, 28% of the global population experienced moderate to severe food insecurity (FAO et al., 2025). This is experienced as having restricted food variety, reduced meal frequency and size, or not having access to food for a day or more (FAO, 2023). Despite food insecurity being defined by limited food access, there is robust evidence linking food insecurity to obesity (Dinour, Bergen & Yeh, 2007; Eskandari et al., 2022; Kowaleski, Wen & Fan, 2018; Metallinos-Katsara, Must, & Gormans, 2012). This relationship is at a greater prevalence in high-income countries and in women (Nettle, Andrews & Bateson, 2017). Moreover, children who experience food insecurity are more likely to be obese than food secure children (Metallinos-Katsara, Must, & Gormans, 2012; Ortiz-Marron et al., 2022; Falcone et al., 2025). Although there is rich correlational data implicating the co-occurrence of food insecurity and obesity, whether food insecurity plays a causative role in the development of obesity and what the driving mechanisms of this relationship are, remain unclear.

Over recent years, the development of rodent models of food insecurity has allowed for these mechanisms to be more closely interrogated. Regardless of the degree of variability within these models (e.g., sex, species, diet type, and food insecurity protocol), these studies show that rodents who experience adolescent/early adulthood food insecurity gain more weight or body fat in adulthood, and this effect is more pronounced in female rodents (Gil et al., 2025; Lin et al., 2022; Myers et al., 2022; Spaulding et al., 2024), thereby demonstrating its predictive validity. These studies also show that food insecurity can lead to altered eating behaviours: female food insecure rodents increase intake when given access to a high fat, high sugar diet (Estacio et al., 2021; Lin et al., 2022), show enhanced food motivation and consume significantly larger meals (Myers et al., 2022). These findings indicate that food insecurity can cause overeating, which could potentially lead to weight gain.

Overeating is driven by the dysregulation of internal physiological (e.g., gut signals) and/or external environmental (e.g., food cues) control of feeding behaviour. However, whether food insecurity leads to changes in the internal and external control of eating has yet to be demonstrated. Evidence from human studies of food insecurity show that children who are food insecure have decreased satiety sensitivity and higher food responsiveness in comparison to food secure children (Kral, Chittams & Moore, 2017; McCurdy et al., 2022), suggesting that food insecurity could affect both internal and external control of feeding. Importantly, deficits in internal and external control of feeding can predict later development of obesity. For example, rats that are less sensitive to the satiating effects of the gut peptide cholecystokinin (CCK) (Wald & Grill., 2019) or exhibit enhanced food cue-induced approach behaviours (Robinson et al., 2014) are more susceptible to diet-induced obesity. Based on these findings, it is possible that food insecurity could dampen internal satiation signalling and increase responsiveness to food cues, leading to overeating and weight gain.

The present study therefore examined the effect of food insecurity on gut signal sensitivity and cue induced appetitive behaviours. Using a rat model of adolescent food insecurity, we showed that previously food insecure rats were less sensitive to the satiating effects of CCK and exhibited enhanced cue-induced appetitive behaviours in a Pavlovian model of appetitive conditioning, when compared to the food secure and food restricted controls. Together, these findings indicate that exposure to food insecurity during adolescence has enduring impacts on feeding behaviours during adulthood, which may lead to subsequent weight gain.

## 2. Methods

### 2.1 Subjects

Juvenile 4-week old female Long Evans rats (UNSW Sydney; N=30) upon arrival were individually housed in ventilated cages within a climate-controlled colony room with a reverse 12hr dark/light cycle (lights on at 7pm). Rats were acclimatised to the colony room for 7 days prior to experimental proceedings, and during this time water and chow (Specialty Feeds, Western Australia) were available *ad libitum*. All experimental procedures were approved by the University of New South Wales Animal Care and Ethics Committee.

### 2.2 Feeding Schedule

The feeding schedule (Figure 1a) began when the rats were 5 weeks of age, prior to which food intake and body weights were monitored daily. Rats were body weight-matched and assigned to either the food secure (n=10) or food insecure (n=10) group. The food secure group had *ad libitum* access to standard chow throughout the study. The food insecure group received a variable amount of chow once daily, at a random timepoint throughout the dark cycle. The amount of chow given ranged from 50-150% of their average daily intake (adapted from Estacio et al., 2021) which was updated weekly based on the food intake of the secure group. Specifically, the average food intake recorded on the final day of the week for the secure group was used as the baseline amount given to the insecure group the following week. We updated the baseline daily intake in this way to account for changes in food intake during adolescence and to achieve similar average weekly chow intake between food insecure and food secure rats. However, insecure rats were still mildly calorie restricted where their average weekly intake was ∼90% of the food secure rats. Consequently, the food restricted group was included to control for the effects of caloric restriction during adolescence. Rats in the food restricted group (n=10) were pair-fed based on the average weekly intake of the insecure group and were fed at a consistent time each day (8am; one hour post dark cycle onset). There were no group differences in body weight and food intake at baseline (Body weight: *F* _(2,29)_ = 1.399, p=0.264; Food intake: *F* _(2,29)_ = 1.285, p=0.293). The feeding schedule ran for 4-weeks, during which water was freely available for all groups, and food intake and body weight were recorded daily.

**Figure 1:**
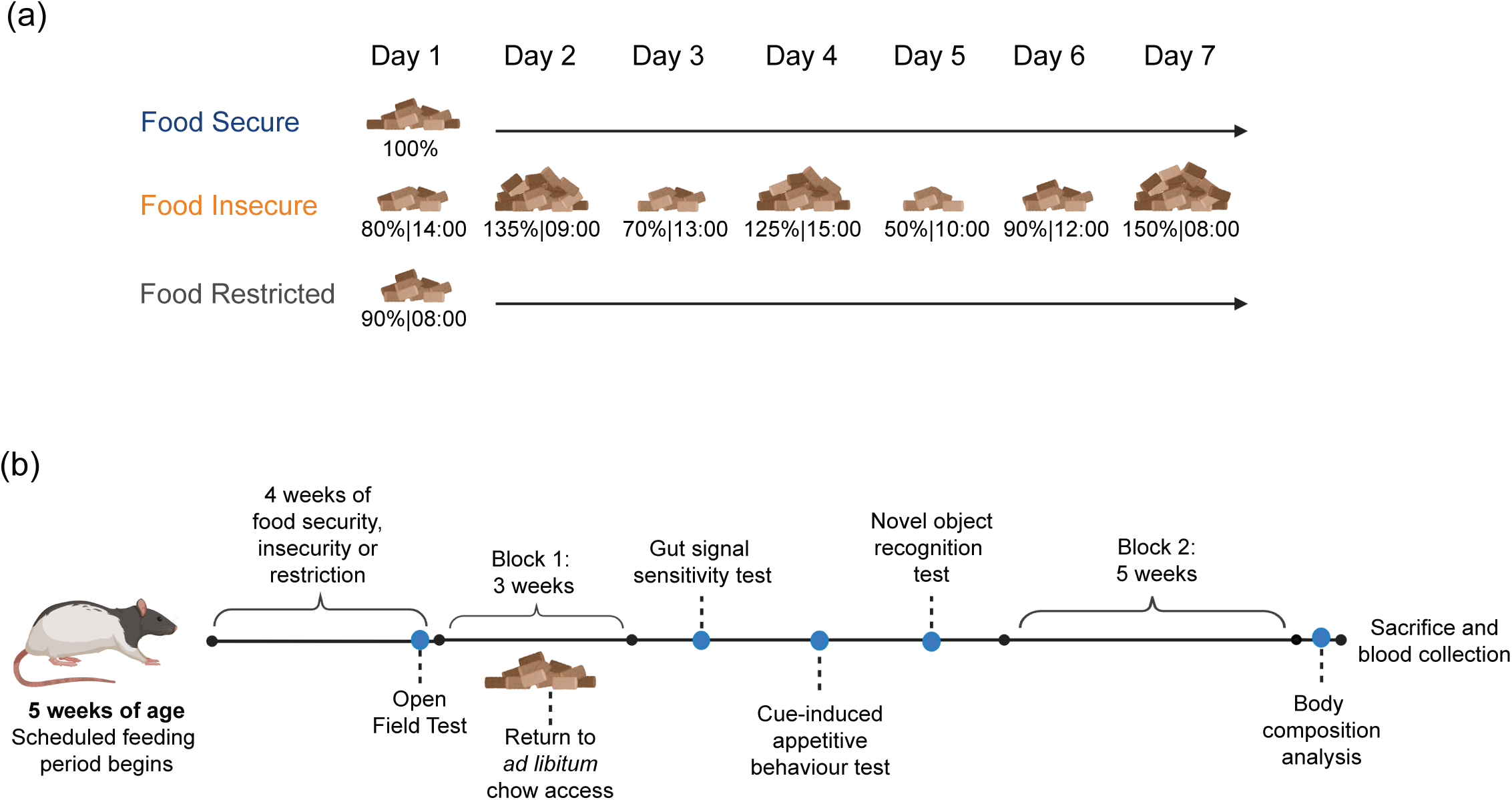
Feeding schedule overview and timeline. (a) Example of the feeding schedule during a given week of the protocol. The amount of chow given is expressed as a percentage of *ad libitum* intake and the time indicates when food is delivered. (b) Female Long Evans rats were assigned to a feeding schedule at 5 weeks of age. The dietary manipulation ran for 4 weeks and concluded with an open field test. Following this, rats returned to an *ad libitum* chow diet for 3 weeks with food intake and body weight monitored (Block 1), before moving onto behavioural testing. The impact of adolescent dietary schedule on feeding behaviours (gut signal sensitivity test, cue-induced appetitive behaviour test), learning and memory (novel object recognition test) was assessed. This was followed by a 5-week *ad libitum* access period (Block 2), where food intake and body weight were monitored in adulthood. At the end of the study, EchoMRI body composition analysis and plasma leptin analysis were performed.

### 2.3 Open Field Test

To determine if food insecurity increases anxiety-like behaviour, an open field test was conducted on the final afternoon of the feeding schedule period (Figure 1b). Rats were placed in the centre of an open field arena (43.2cm x 43.2cm x 30.5cm) and were allowed to explore for 30 minutes. The open field arena consisted of 16×16 rows of photocell beams to allow activity detection and registration by a connected computer system. Additionally, the arena was divided into 2 zones, whereby the centre area was classified as the ‘middle’ and the surrounding area was classified as ‘outer’. Distance travelled and time spent active across each zone was recorded, and percent activity in the middle zone relative to total activity [middle zone activity/ total activity across both zones] x 100 was calculated as a measure of anxiety-like behaviour.

### 2.4 Food intake and body weight

After 4 weeks on their respective feeding schedule, all groups had *ad libitum* access to chow for the remainder of the experiment. This selectively isolated the experience of food insecurity and restriction to the adolescent window. To determine whether food insecurity results in metabolic changes after switching to an *ad libitum* chow diet, food intake and body weight were recorded for the first 5 days of *ad libitum* access and weekly for 3 weeks prior to feeding behaviour tests (Block 1), and 5 weeks after feeding behaviour tests (Block 2) (Figure 1b).

### 2.5 Gut Signal Sensitivity Test

To assess gut signal sensitivity, we examined the effects of the gut peptide CCK on intake of a 10% sucrose solution. Rats were given 30-minute access to 10% sucrose in a bottle connected to a lickometer in a standard Med Associates chamber. Each lick broke a photobeam that recorded the number of licks. All rats were habituated to consuming sucrose in the chamber and to receiving intraperitoneal injections of saline (0.9% sodium chloride) for 3 days. During test days, rats received 1mL/kg intraperitoneal injections of either vehicle (0.9% sodium chloride), 1.5 µg/kg CCK or 3 µg/kg CCK. Immediately after injections, rats were placed into the chamber and licking behaviour across the session was recorded. Testing was conducted in a within subjects counterbalanced design and each test day was separated by at least a 24-hour washout period. Total licks after 15 minutes and 30 minutes were recorded and lick microstructure analysis was conducted. A burst is defined as a series of minimum three licks with an inter-lick interval of < 500ms (Johnson, 2018). Burst analysis was conducted on vehicle test days.

### 2.6 Cue-Induced Appetitive Behaviour

To assess cue-induced appetitive behaviour, we used a Pavlovian appetitive conditioning task where rats learnt to associate a cue with a food reward. Rats underwent 70-minute daily training sessions over a period of 10 days in standard Med Associates chambers. These sessions consisted of eight 15 seconds presentations of an auditory cue (CS+) that co-terminated with the delivery of two sucrose pellets, and eight 15 seconds presentations of a different auditory cue (CS-) that was non-reinforced. The cue types (tone and clicker) were counterbalanced across rats such that half the rats received the tone as CS+, clicker as CS-and the other half received clicker as CS+, tone as CS-. The CS+ and CS-trials were presented in a randomised order with an intertrial interval of 4 minutes. On test day, rats underwent the same Pavlovian appetitive conditioning task but food reward was not delivered following CS+ presentation. Magazine entries before, during and after each CS presentation were recorded. Appetitive behaviour was quantified as ‘Elevation score’ which is the difference in magazine entries 15 seconds before and 15 seconds during CS presentation. Additionally, latency to enter the magazine following CS+ onset was manually scored from video recordings on the final day of training.

### 2.7 Novel Object Recognition Test

To ensure that any effects observed with feeding behaviours were not generalised to differences in learning and memory, we conducted a novel object recognition test. Testing took place in the same open field arena as previously mentioned, and a video camera was placed above the boxes to video record exploration behaviour. Three different shaped objects were used in this task. During the familiarisation session, Object 1 and Object 2 were placed in the centre of the open field box an equidistance apart, and rats were allowed to explore freely for 10 minutes. Following a 24-hour retention period, Object 1 was swapped out for Object 3 and rats were allowed to explore for 10 minutes. Exploratory behaviour was defined as nose-directed behaviour towards the objects (i.e. sniffing). Climbing onto or chewing the object was not considered exploratory behaviour (Leger et al., 2013). To quantify whether rats showed a novelty preference, the time spent exploring the novel object was divided by the total exploration time of both objects at test (Tran & Westbrook, 2018).

### 2.8 Body Composition

To determine if adolescent food insecurity in this study led to long-term changes in body fat mass and lean mass, we measured body composition using EchoMRI at UNSW Biological Resources Imaging Laboratory. Percent fat mass and percent lean mass were calculated as [fat mass (g)/ body weight (g)] x 100 and [lean mass(g)/ body weight (g)] x 100.

### 2.9 Plasma Leptin

A subset of rats from each group (n=5/group) were sacrificed with an injection of lethabarb and blood collected following a snip of the descending aorta. Blood was collected in EDTA tubes on ice, centrifuged at 4000 rpm for 15 minutes at 4°C. Plasma was extracted, aliquoted into tubes and stored at -20°C until further analysis. Plasma leptin levels were measured using Rat Leptin ELISA kit (RAB0335-1KT, Millipore) according to manufacturer’s instructions.

### 2.10 Data Analysis

Statistical analyses were conducted using IBM SPSS Statistics and figures were produced using Prism software. Data were presented as mean **±** standard error of the mean (SEM) in all figures and statistical significance was considered at p<0.05. Data from the feeding periods and cue-induced feeding test were analysed using mixed analysis of variance (ANOVA) with Tukey post hoc tests when significance was observed. Data from the open field test, novel object recognition test, EchoMRI and plasma leptin were analysed using a one-way ANOVA with Tukey post hoc tests when significance was observed. Data from the gut signal sensitivity test were analysed using a repeated measures ANOVA with Bonferroni pair-wise comparisons when significance was observed. One rat in the food restricted group died unexpectedly at the start of the feeding behaviour test, resulting in a final sample size of n=9 food restricted rats for the remainder of the study.

## 3. Results

### 3.1 Effects of feeding schedule on food intake and body weight during the 4-week schedule period

Repeated measures ANOVA on weekly average food intake revealed a significant main effect of time (*F* _(2,57)_ = 49.702, p<0.001), group (*F*_(2,27)_ = 9.474, p<0.001), but no significant time x group interaction (*F*_(4,57)_ = 1.932, p=0.113) (Figure 2a). This suggests that overall food intake differed between the groups, however the pattern of change in food intake was the same for all groups. Tukey post hoc test revealed that the food secure rats ate significantly more than the food restricted (p<0.001) and food insecure (p<0.01) rats. This makes sense, given that the food insecure and restricted groups were consuming 90% of the average weekly intake of the food secure rats.

**Figure 2:**
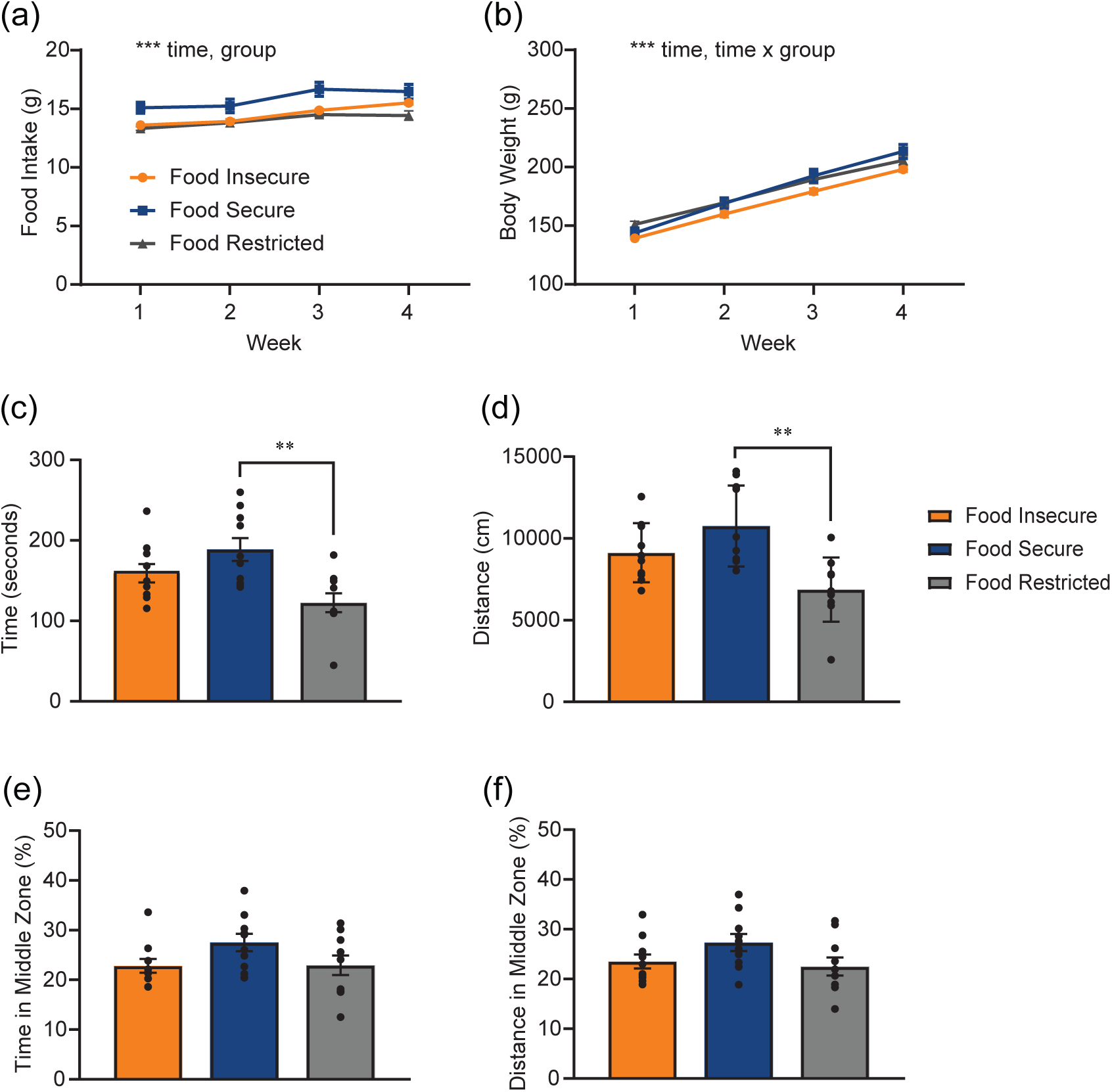
Food intake, body weight and anxiety-like behaviours during adolescent feeding schedule. Food secure rats showed higher (a) weekly food intake compared to insecure and restricted groups. Whilst there was a time x group interaction for (b) body weight, there were no group differences at any week. In the open field test, food restricted rats showed reduced (c) total active time and (d) total distance travelled when compared to food secure rats. There were no group differences in (e) the percent time spent in middle zone and (f) the percent distance travelled in middle zone. ** p<0.01; *** p<0.001.

For body weight, a mixed ANOVA revealed a significant main effect of time (*F*_(1,38)_ = 2792.813, p<0.001) and time x group interaction (*F*_(3,38)_ = 12.692, p<0.001) but no main effect of group (*F*_(2,27)_ = 2.291, p=0.12). Follow up one-way ANOVA found no differences in average body weight between groups across any of the weeks (p>0.05) (Figure 2b). This indicates that although there was no consistent overall difference in body weight between groups, changes in body weight across the 4-week period differed depending on diet schedule.

### 3.2 Effects of feeding schedule on anxiety-like behaviours

The open field test was used to measure anxiety-like behaviours on the final day of the feeding schedule period. We first investigated whether there were differences in overall distance travelled and time spent moving (active time) between groups. We observed significant group differences in both distance travelled (one-way ANOVA: *F*_(2,27)_ = 7.081, p=0.003) and active time (*F*_(2,27)_ = 8.652, p=0.001). Post hoc Tukey tests indicate that the food restricted group spent significantly less time moving (p=0.002) and travelled less distance (p=0.001) than the food secure group, although the food insecure group was not different to either group on both measures (p>0.05) (Figure 2c,d). To determine anxiety-like behaviour, we compared percent activity in the middle zone relative to total activity across groups. We observed no group differences on percent active time in middle zone (*F*_(2,27)_ = 2.95, p=0.069), or percent distance travelled in middle zone (*F*_(2,27)_ = 2.347, p=0.115) (Figure 2 e, f). Taken together, these results suggest that whilst food restricted rats showed lower overall locomotor activity compared to food secure rats, the lack of difference in middle zone activity suggests no differences in anxiety-like behaviours between groups.

### 3.3 Effects of feeding schedule on food intake and body weight prior to feeding behaviour testing

Following the 4-week feeding schedule, all groups were given *ad libitum* access to chow. To measure the acute effects of switching to an ad-libitum diet, food intake and body weight were monitored for 3 weeks: daily for the first 5 days, and weekly thereafter.

#### 3.3.1 Day 1 – 5 of *ad libitum* chow access

A mixed ANOVA on food intake during the first 5 days showed a significant main effect of time (*F*_(4,108)_ = 4.709, p=0.002) and time x group interaction (*F*_(4,108)_ = 5.232, p<0.001) but no group effect (*F*_(2,27)_ = 1.514, p=0.238). This indicates that changes in food intake across this 5-day period differed between the groups, however their overall intake is similar. Follow-up one-way ANOVA revealed no differences in food intake between groups on Day 1, 4 or 5 but there were statistically significant differences on Day 2 (*F*_(2,27)_ = 5.69, p=0.009) and Day 3 (*F*_(2,27)_ = 4.405, p=0.022). Post hoc Tukey tests indicate that on Day 2 the food restricted group ate significantly less than both the food insecure (p=0.045) and secure group (p=0.01), and this effect continued into Day 3 (p<0.05) (Figure 3a).

**Figure 3:**
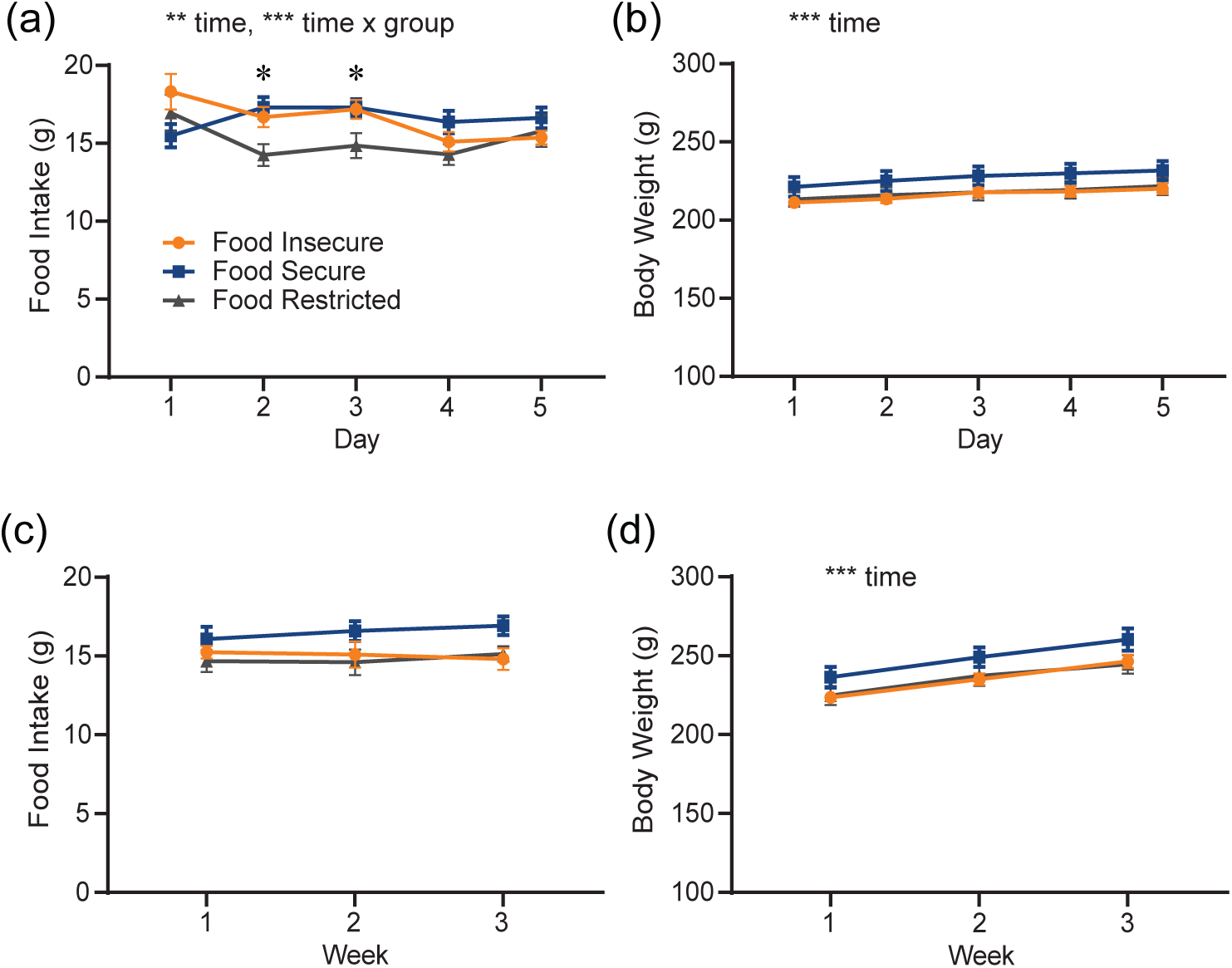
Impact of adolescent dietary schedule on food intake and body weight following return to ad libitum chow access. During the first 5 days of Block 1, food restricted rats had lower (a) daily food intake during Days 2 and 3 when compared to secure and insecure rats. There were no effects on (b) body weight. During the three weeks in Block 1, there were no group differences on (c) weekly food intake and (d) body weight. * p<0.05; ** p<0.01; *** p<0.001.

For body weight, we did not observe any acute group differences during the first 5 days (mixed ANOVA: *F*_(2,27)_ = 1.458, p=0.25). There was a significant main effect of time (*F*_(4,108)_ = 80.838, p<0.001) (Figure 3b) indicating that body weight gain during this period was consistent for all groups.

#### 3.3.2 Week 1 – 3 of *ad libitum* chow access

Next, we analysed these measures across 3 weeks prior to behaviour testing using mixed ANOVA. Food intake was not affected by time (*F* _(2,54)_ = 0.267, p=0.767) or previous diet schedule (*F* _(2,27)_ = 3.03, p=0.066) (Figure 3c). Body weight increased consistently for all groups during this period (main effect of time: *F* _(2,54)_ = 246.397, p<0.001), although there was no effect of group (*F* _(2,37)_ = 1.944, p=0.163) (Figure 3d). Taken together, prior feeding schedule did not impact food intake or body weight during this period.

### 3.4 Effects of food insecurity on gut signal sensitivity

During the gut signal sensitivity test, the effects of vehicle, 1.5 µg/kg, and 3 µg/kg CCK on 10% sucrose intake were measured by quantifying total licks at 15 minutes and 30 minutes. To assess sensitivity to CCK within each group, repeated measures ANOVA were performed at each time point. There was a main effect of treatment for the food insecure (*F*_(2,18)_ = 6.42, p=0.008), food secure (*F*_(2,18)_ = 6.138, p=0.009) and food restricted (*F*_(2,16)_ = 13.075, p<0.001) groups. Follow-up Bonferroni pairwise comparisons found that food insecure rats licked significantly less following 3 µg/kg CCK (p=0.027), but not 1.5 µg/kg CCK injection compared to vehicle (p>0.05). The food secure rats licked significantly less following 1.5µg/kg CCK (p=0.029), with a trend towards significant reduction in licks when injected with 3µg/kg CCK (p=0.075) compared to vehicle. The food restricted group significantly reduced licking following both 1.5µg/kg CCK (p=0.039) and 3µg/kg CCK (p<0.001) injection, relative to vehicle (Figure 4b). At 30 minutes however, there was no effect of treatment on number of licks for the previously food insecure (*F*_(2,18)_ =2.223, p=0.137), food secure (*F*_(2,18)_ =1.598, p=0.23) and restricted rats (*F*_(2,16)_ = 2.906, p=0.084) (Figure 4c). Overall, whilst all groups reduced sucrose licks following CCK administration, only the food secure and food restricted groups reduced the number of licks at the lower dose of CCK. This suggests that the food insecure group showed less sensitivity to the satiating effects of CCK.

**Figure 4:**
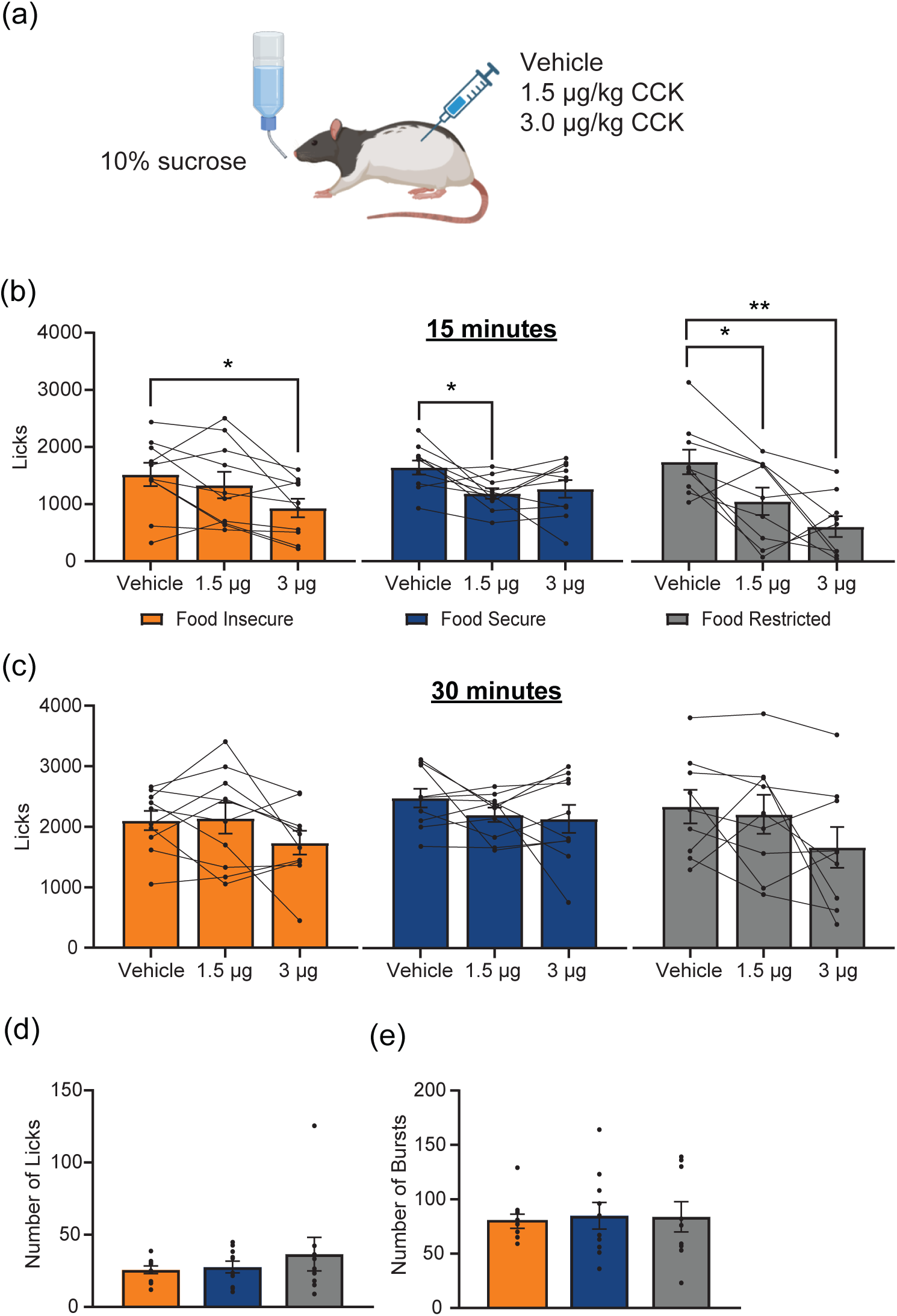
Adolescent food insecurity dampened gut signal sensitivity. (a) Schematic of experimental design: Rats had access to a 10% sucrose solution and licking behaviour was recorded using a lickometer. Across test days, rats were injected with either vehicle,1.5 μg/kg CCK or 3.0 μg/kg CCK, with a 24-hour washout period in between. On test days licking behaviour was analysed following (b) 15 minutes and (c) 30 minutes for each group. (b) At 15 minutes, CCK significantly reduced the number of licks across all groups, where the food restricted and food secure groups suppressed intake to 1.5 μg/kg CCK, whereas the food insecure only suppressed intake following 3.0 μg/kg CCK. (c) At 30 minutes, there was no effect of CCK on licking behaviour across groups. There were no group differences in the (d) average number of licks per burst or (e) the number of bursts initiated. * p<0.05; ** p<0.01.

We also assessed their lick microstructure by quantifying the average burst size and number for each group to determine whether there were any differences in palatability. Data were analysed using one-way ANOVA. There was no effect of group on the average size of each burst (*F*_(2,28)_ =0.685, p=0.513) or the number of bursts initiated (*F*_(2,28)_ =0.6, p=0.942) (Figure 4 f, e). This indicates that there were no differences in the lick microstructure and palatability between groups.

### 3.5 Effects of food insecurity on cue induced appetitive behaviours

To investigate the effect of adolescent dietary schedule on cue-induced appetitive behaviours, we analysed their appetitive responses to a CS during training and at test. One rat from the food restricted group was removed from this analysis as it was a statistical outlier.

Rats learnt to differentially respond to the CSs across training days, showing greater elevation scores to the CS+ compared to the CS-, as confirmed by a time x CS interaction (mixed ANOVA: *F*_(4,110)_ = 168.95, p<0.001). Additionally, there was a main effect of group (*F*_(2,26)_ = 7.316, p=0.003) and a significant CS x group interaction (*F*_(2,260)_= 4.025, p=0.03), indicating that responses to the CS+ and CS-during training were influenced by adolescent feeding schedule. Post hoc Tukey test revealed that CS+ elevation score was significantly higher in food insecure group compared to the food secure (p=0.046) and restricted groups (p=0.003).

This suggests that prior food insecurity increased overall responding to the food-associated cue during training (Figure 5b). We were then interested to determine the point at which groups started to discriminatively respond to the cues during training. Paired samples t-tests comparing responding to the CS+ versus CS-within each group revealed that the food insecure discriminatively responded to the cues from Day 1 (t(9) = 3.125, p=0.012), Day 2 for the food restricted group (t(8) = 2.747, p=0.025) and Day 4 for the food secure (t(9) = 4.206, p=0.002) of training (Figure 5b). Hence, it appears that the food insecure and restricted groups learnt the food-cue association earlier than the food secure group. Additionally, we also measured latency to enter the magazine following CS+ onset on the last day of training. Unfortunately, due to a video recording error, we only have data from the food insecure and secure groups for this measure, and hence an independent samples t-test was performed to compare group differences. We found that the food insecure rats had a significantly shorter latency to enter the magazine following CS+ onset compared to the food secure rats (*t*(18) = - 3.635, p= 0.002) (Figure 5c). This suggests the food insecure rats had greater behavioural activation to the food-cue compared to food secure rats. At test, appetitive behaviours were measured in the absence of food reward. There was a significant main effect of CS (*F*_(1,26)_ = 121.78, p <0.001), group (*F*_(2,26)_= 5.13, p=0.013) and CS x group interaction (*F*_(2,26)_ = 8.51, p=0.001). Tukey post hoc tests reveal that the food insecure group showed significantly higher appetitive behaviours compared the food secure (p=0.04) and food restricted (p=0.019) at test (Figure 5d). This suggests that adolescent food insecurity increased responding to the food-associated cue.

**Figure 5:**
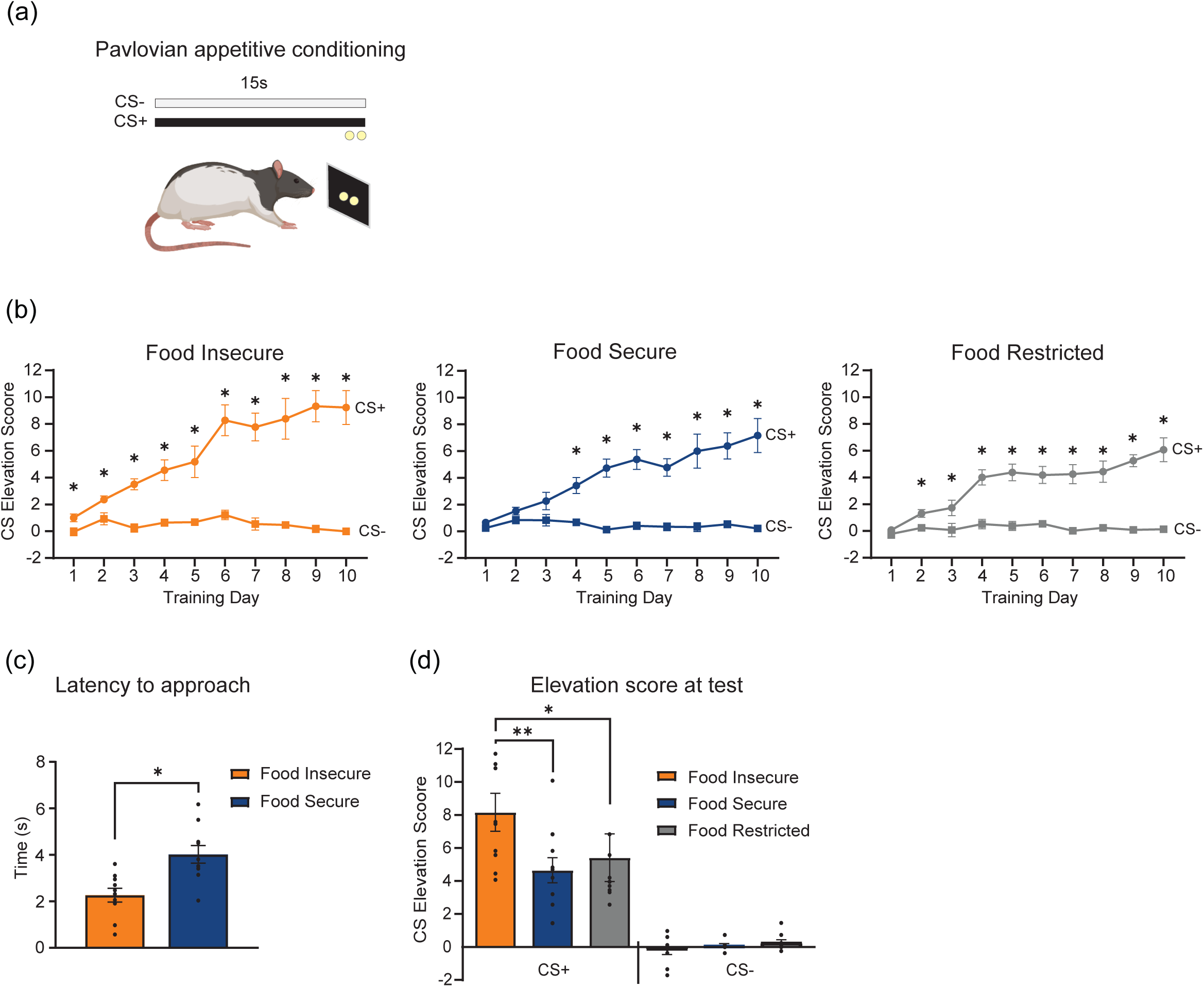
Adolescent food insecurity enhanced acquisition and expression of cue induced appetitive behaviours. (a) Schematic of experimental design: During training, rats were presented with two auditory cues; one that co-terminated with the delivery of two sucrose pellets (CS+) and another that was unreinforced (CS-). Rats learnt to associate the CS+ with reward delivery and subsequently elicited appetitive behaviours (magazine entries). At test, cue-induced appetitive behaviours were recorded in the absence of reward. (b) All groups showed discriminative responding to CS+ and CS-during training. Elevation scores for the CS+ and CS-were significantly different after 1 day (Insecure), 4 days (Secure) and 2 days (Restricted) of training. (c) The food insecure group had reduced latency to approach the magazine following CS+ onset when compared to the food secure group. (d) At test, the food insecure group showed elevated CS+ appetitive behaviours compared to secure and restricted groups. * p<0.05; ** p<0.01.

### 3.6 Effects of feeding schedule on learning and memory

Given that food insecure rats learnt to discriminate between food paired CS+ vs CS-faster than food secure and restricted groups, we wanted to investigate whether food insecure rats have a generalised increase in learning and memory. To do this, we conducted a novel object recognition test. A one-way ANOVA showed no significant difference in novelty preference scores between groups (*F*_(2,27)_ = 1.817, p=0.182) (Figure 6b). This indicates that there is no difference in learning and memory between groups.

**Figure 6:**
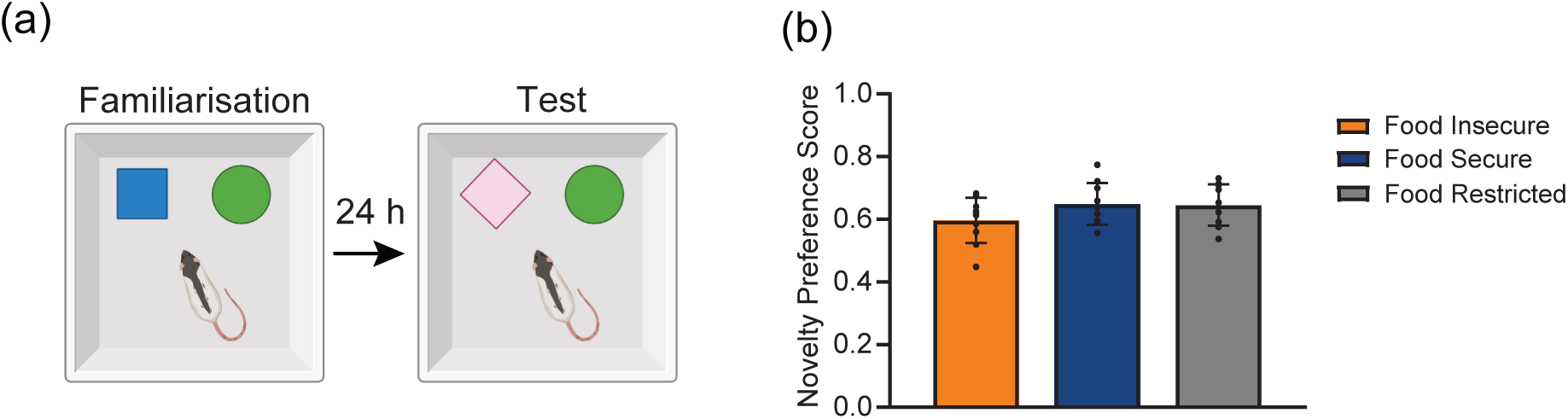
Adolescent feeding schedule did not impact learning and memory. (a) Schematic of novel object recognition test: Rats were allowed to freely explore 2 objects for two 10-minute sessions separated by a 24-hour retention period. At test, a familiar object was swapped out for a novel object, and subsequent exploration behaviours were measured. (b) Adolescent feeding schedule had no effect on novelty preference scores during the novel object recognition test.

### 3.7 Effects of food insecurity on food intake and body weight after feeding behaviour testing

Following feeding behaviour testing, body weight and food intake were measured for 5 weeks. We observed a main effect of time (*F*_(4,104)_ = 5.752, p<0.001) and group (*F*_(2,26)_ = 7.295, p=0.03) on food intake during this period, indicating that food intake differed depending on adolescent feeding schedule. Post hoc Tukey tests reveal that the restricted group ate significantly less than both the previously insecure (p=0.013) and secure (p=0.004) groups, although there was no difference between the secure and insecure groups (p>0.05) (Figure 7a). This indicates that adolescent caloric restriction may reduce food intake in adulthood.

**Figure 7:**
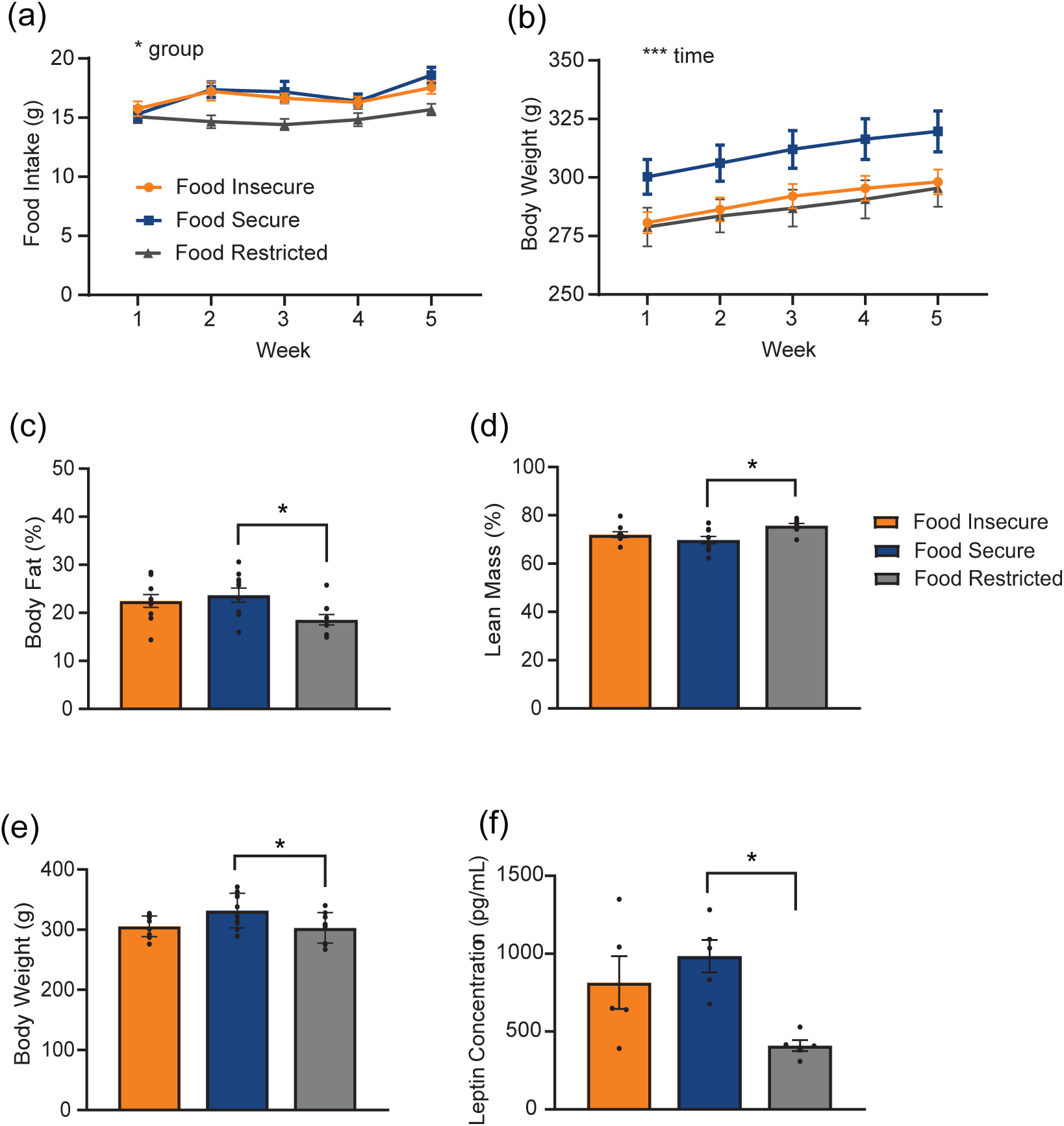
Impact of adolescent dietary schedule on body weight, food intake and body composition during adulthood. Food restricted rats showed (a) lower weekly food intake than secure and insecure rats, but not (b) weekly body weight. Compared to food secure rats, food restricted rats had (c) lower percent body fat, (d) higher percent lean mass, (e) lower adult body weight and (f) lower plasma leptin levels. Food insecure rats were not different from secure or restricted rats in (c) – (f). * p<0.05; ** p<0.01; *** p<0.001.

We also observed that across the 5-week period, body weight increased for all groups (*F*_(3, 75)_ = 141.532, p<0.001) and that there was a trend towards a significant effect of adolescent food schedule on body weight (*F*_(3,75)_ = 3.346, p=0.051) (Figure 7b).

### 3.8 Effects of food insecurity on body composition during adulthood

To determine the impact of adolescent feeding schedule on body composition, rats were placed in the EchoMRI instrument. Body weight, percent lean mass and percent fat mass were analysed using one way ANOVA. We found that adolescent feeding schedule significantly affected percent fat mass (*F*_(2,26)_ = 4.027, p=0.03) and percent lean mass (*F*_(2,26)_ = 5.673, p=0.009) in adulthood. Post hoc Tukey test revealed that the food restricted group had significantly less percent fat mass compared to the food secure group (p=0.029), however the food insecure group was not significantly different to either group (p>0.05) (Figure 7c).

Similarly, the food restricted group had significantly more percent lean mass compared to the food secure group (p=0.007), however the food insecure rats was not significantly different to either group (p>0.05) (Figure 7d). At the final weigh-in, we found that adolescent dietary schedule significantly impacted body weight (*F*_(2,26)_ = 4.226, p=0.026). Post hoc Tukey tests revealed that the food restricted rats weighed significantly less than the food secure rats (p=0.04), however the food insecure rats were not significantly different to either group (p>0.05) (Figure 7e). This suggests that adolescent caloric restriction can impact body weight in adulthood. Taken together, adolescent caloric restriction decreased body weight through reduced fat mass in adulthood, although adolescent food insecurity somewhat mitigated these effects.

### 3.9 Effects of food insecurity on plasma leptin levels

One way ANOVA analysis on plasma leptin levels revealed a main effect of group (*F*_(2,14)_ = 6.369, p=0.013). Tukey’s post hoc test further showed that plasma leptin levels in food restricted rats were significantly lower compared to food secure rats (p=0.012), with a trend towards a significant reduction when compared to food insecure rats (p=0.073). Plasma leptin levels were not different between food secure and food insecure rats (p>0.05) (Figure 7f). Similar to the effects on body composition, adolescent caloric restriction resulted in lower plasma leptin levels that were partially mitigated by adolescent food insecurity.

## 4. Discussion

In the present study, we investigated the effects of food insecurity on gut signal sensitivity and cue-induced feeding behaviours. Using a rodent model of food insecurity, where unpredictability was maximised by varying the time of food delivery and quantity given each day, we conducted a series of behavioural tests measuring feeding behaviours, learning, memory and anxiety-like behaviour. Although adolescent feeding schedule did not impact memory and anxiety-like behaviour, food insecurity dampened the satiating effects of CCK and enhanced the acquisition and expression of an appetitive response to a food-associated cue. Notably, these effects were observed at least 3 weeks after the scheduled feeding period and prior to any differences in body weight. This indicates that food insecurity during adolescence can produce adaptations that promote increased food intake in adulthood, even when access to food has been restored. This was reflected in our finding that the food insecure rats ate significantly more than the food restricted rats during adulthood. Moreover, body composition in adulthood was significantly impacted by feeding schedule. Food restricted rats showed greater lean mass, reduced fat mass and weighed significantly less than food secure rats, however food insecure rats did not differ from either group. This suggests that these effects of food restriction on body composition can be somewhat mitigated when restriction is coupled with food insecurity. Taken together, these findings demonstrate that experiencing food insecurity during adolescence can lead to enduring metabolic and behavioural adaptations that may serve as mechanisms through which food insecurity induces obesity.

One of our key findings showed that previously food insecure rats were less sensitive to the satiating effects of the gut satiation signal CCK. When injected with a lower dose of CCK (1.5µg/kg), both the food secure and restricted groups reduced intake, whereas this dose had no effect on the food insecure rats. This demonstrates that having unpredictable access to food during adolescence can result in impaired sensitivity to gut satiation signals later in life. This finding is also consistent with a previous study which showed that food insecure rats consumed more and had larger meals compared to food secure rats during an 8-hour access period to a palatable liquid diet (Myers et al., 2022), which suggests reduced satiation. This pattern of ‘overeating’ has similarly been observed in human studies. Food insecure participants overeat in comparison to food secure participants when food is available *ad libitum*, even if recent intake has been equated (Nettle et al., 2019; Stinson et al., 2018; Rasmusson et al., 2019). Children experiencing food insecurity have been found to eat past perceived fullness and in absence of hunger (McCurdy et al., 2022). Therefore, it is possible that food insecurity reduces sensitivity to gut signalling, which may lead to excessive food intake.

That food insecurity, but not caloric restriction, dampens the satiating effects of CCK suggests that variability in food access can have profound impact on gut signal processing. Indeed, studies in humans manipulating daily meal frequency showed that irregular meal frequency reduces subjective fullness and dietary thermogenesis (increased metabolic rate following food intake) when compared to individuals with regular meal frequency (Alhussain, Macdonald & Taylor, 2016, 2022; Farshchi, Taylor & MacDonald, 2004). In contrast, intermittent fasting - a dietary pattern that is defined by consistent long periods of deprivation between meals increases fullness and improves metabolic outcomes, findings that are opposite to that of food insecurity (Chaix et al., 2014; Varady et al., 2022; Welton et al., 2020). This demonstrates that simply having a variable pattern of eating, rather than caloric restriction, can cause metabolic adaptations that favour a positive energy balance (Alhussain, Macdonald & Taylor, 2022).

The mechanisms underlying the impact of food insecurity on gut signal sensitivity are still unclear but current theories suggest that unpredictability of food access can inhibit the body’s ability to appropriately adapt and prepare for food intake, by limiting the appropriate digestive and metabolic anticipatory responses that are necessary for regulating food intake (cephalic responses) (Power & Schulkin, 2008; Skvortsova et al., 2021; Smeets, Erkner & Graaf, 2010). This can affect subsequent satiation signalling (Smeets, Erkner & Graaf, 2010) and their associated neural responses, particularly within the nucleus of the solitary tract (NTS) as rats with reduced responsiveness to CCK also show reduced CCK-induced activation of NTS neurons (Wald and Grill, 2019). Thus, future research should examine whether food insecurity affects cephalic and satiation signalling and NTS neural responses associated with impaired gut signal sensitivity.

In addition to impairments in gut signal sensitivity, the current study was the first to show that previously food insecure rats had enhanced learning of a food-associated cue, greater anticipatory response to the cue and elevated cue-induced appetitive behaviours. This shows that food insecurity can heighten responses to food-associated cues. This is supported by existing theories which suggest that unpredictability magnifies the appetitive properties of the reward, leading to significantly enhanced motivational salience of the reward and reward-related cues (Anselme, 2013; Anselme & Güntürkün, 2019; Robinison et al., 2014). Importantly this adaptation to reward uncertainty may have evolutionary significance, as increased motivation to food and food-related cues would enhance the likelihood of survival when resources are scarce (Anselme & Güntürkün, 2019; Robinson, Bonmariage & Samaha, 2023). The role of unpredictability in driving cue-induced approach behaviours is further supported by a recent study which introduced unpredictability into an intermittent access model, such that the length of access and amount of sucrose solution available varied each session. The study showed that rats subjected to unpredictable intermittent access to sucrose exhibited heightened responding to the CS+ and increased CS-reinstatement behaviour (Robinson, Bonmariage & Samaha, 2023). Therefore, uncertainty itself can be a key driver of food-motivated behaviours.

There is evidence that food uncertainty activates and sensitises dopamine signalling, and in turn drives reward-seeking behaviour (Anselme, 2013; Anselme & Güntürkün, 2019; Fiorillo, Tobler & Schultz, 2003). It is possible that experiencing food insecurity during a critical period of development (adolescence) may sensitise dopamine neurons to the rewarding properties of food, and have lasting impacts on food-motivated behaviours, such as responding to a food-associated cue. Thus far, only one study has demonstrated changes in striatal dopamine signalling following adolescent food insecurity (Lin et al., 2022), though this was shown only in male rats and was unrelated to the impact of food insecurity on feeding behaviours during adulthood. Thus, future research elucidating the neurobiological mechanisms of adolescent food insecurity on cue-induced feeding behaviours is warranted.

The central aim of our study was to determine whether adolescent food insecurity causes dysregulated feeding behaviours that promote the development of obesity. Although we showed that adolescent food insecurity led to dysregulated feeding behaviours, it did not increase weight gain in adulthood. At the final weigh in and during the 5-weeks post-behaviour testing, the body weight of food insecure rats was not different to either the food restricted or food secure group, although food restricted rats weighed significantly less than food secure rats. A similar pattern of result was observed when comparing percent fat mass, lean mass and plasma leptin levels. These findings indicate that adolescent caloric restriction has a significant impact on body composition and weight in adulthood. Given that food insecure rats were not different to either group, this suggest that food insecurity partially attenuated the effects of caloric restriction, and that adaptations to the unpredictability of food access, rather than overall caloric intake are driving these outcomes. Interestingly, the food restricted group had significantly lower food intake compared to both other groups in the 5 weeks following behaviour testing. This indicates that although both groups experienced the same adolescent caloric restriction, food insecurity promoted greater food intake behaviours and weight gain during adulthood. Previous research showed that compared to rats with *ad libitum* access, rats who experience caloric restriction during adolescence do not gain as much weight after access to food is restored (Brownlow et al., 1993; Hubert et al., 2000). It is likely that food restriction during this critical period delays the onset of puberty and causes growth retardation, given that puberty onset is intertwined with energy balance systems (Holehan & Merry, 1985; Rizzoto et al., 2019). However, when caloric restriction was coupled with unpredictability during adolescence, this effect was attenuated. This demonstrates the significant impact that unpredictability has on subsequent body weight. Exploring how different severities of food insecurity can shape later food intake behaviours and body weight using rodent models should be targeted in future research.

In conclusion, we found that unpredictable quantity and time of food delivery confined to the adolescent window significantly reduced gut signal sensitivity and heightened cue-induced feeding behaviours. Importantly, we were able to show these differences occurred prior to any differences in weight and many weeks after our feeding schedule, highlighting the enduring metabolic and behavioural impact of food insecurity. The developmental impact of food insecurity on subsequent food intake behaviours may be driven by neural underpinnings, and future mechanistic research should be aimed at uncovering these to contribute to a better understanding of the food insecurity-obesity paradox.

## Acknowledgements

Authors would like to thank Prof Belinda Liddell for early conceptual discussions that motivated this study and for feedback on the manuscript. Authors would also like to thank Jo Ann Yap for assisting with the plasma leptin ELISA experiment and analysis.

## Funding sources

This work was supported by the Australian Research Council FT220100711 (ZYO).

## Data availability

Data will be made available upon request.

## Notes

### Competing Interest Statement

The authors have declared no competing interest.

## References

Alhussain, M. H., Macdonald, I. A., & Taylor, M. A. (2016). Irregular meal-pattern effects on energy expenditure, metabolism, and appetite regulation: a randomized controlled trial in healthy normal-weight women. The American journal of clinical nutrition, 104(1), 21–32. 10.3945/ajcn.115.125401

Alhussain, M. H., Macdonald, I. A., & Taylor, M. A. (2022). Impact of isoenergetic intake of irregular meal patterns on thermogenesis, glucose metabolism, and appetite: a randomized controlled trial. The American Journal of Clinical Nutrition, 115(1), 284–297. 10.1093/ajcn/nqab323

Anselme, P. (2013). Dopamine, motivation, and the evolutionary significance of gambling-like behaviour. Behavioural brain research, 256, 1–4. 10.1016/j.bbr.2013.07.039

Anselme, P., & Güntürkün, O. (2019). How foraging works: uncertainty magnifies food-seeking motivation. Behavioral and Brain Sciences, 42, e35. doi:10.1017/S0140525X18000948

Brownlow, B. S., Park, C. R., Schwartz, R. S., & Woods, S. C. (1993). Effect of meal pattern during food restriction on body weight loss and recovery after refeeding. Physiology & behavior, 53(3), 421–424. 10.1016/0031-9384(93)90133-Z

Chaix, A., Zarrinpar, A., Miu, P., & Panda, S. (2014). Time-restricted feeding is a preventative and therapeutic intervention against diverse nutritional challenges. Cell metabolism, 20(6), 991–1005. 10.1016/j.cmet.2014.11.001

Dinour, L. M., Bergen, D., & Yeh, M. C. (2007). The food insecurity–obesity paradox: a review of the literature and the role food stamps may play. Journal of the American Dietetic Association, 107(11), 1952–1961. 10.1016/j.jada.2007.08.006

Eskandari, F., Lake, A. A., Rose, K., Butler, M., & O’Malley, C. (2022). A mixedLmethod systematic review and metaLanalysis of the influences of food environments and food insecurity on obesity in highLincome countries. Food science & nutrition, 10(11), 3689–3723. 10.1002/fsn3.2969

Estacio, S. M., Thursby, M. M., Simms, N. C., Orozco, V. A., Wu, J. P., Miawotoe, A. A., Worth, W.W., Capeloto, C.B., Yamashita, K., Tewahade, K.R., & Saxton, K. B. (2021). Food insecurity in older female mice affects food consumption, coping behaviors, and memory. Plos one, 16(4), e0250585. 10.1371/journal.pone.0250585

Farshchi, H. R., Taylor, M. A., & Macdonald, I. A. (2004). Decreased thermic effect of food after an irregular compared with a regular meal pattern in healthy lean women. International Journal of Obesity, 28(5), 653–660. 10.1038/sj.ijo.0802616

Fiorillo, C. D., Tobler, P. N., & Schultz, W. (2003). Discrete coding of reward probability and uncertainty by dopamine neurons. Science, 299(5614), 1898–1902. 10.1126/science.1077349

Food and Agriculture Organization of the United Nations. (2023). Hunger and food insecurity. Retrieved from: https://www.fao.org/hunger/en/

FAO, IFAD, UNICEF, WFP and WHO. (2025). The State of Food Security and Nutrition in the World 2025 – Addressing high food price inflation for food security and nutrition. 10.4060/cd6008en

Gil, C. R., Lund, J., Żylicz, J. J., RaneaLRobles, P., Sørensen, T. I., & Clemmensen, C. (2025). Food insecurity promotes adiposity in mice. Obesity, 33(6), 1087–1100. 10.1002/oby.24259Digital Object Identifier (DOI)

Holehan, A. M., & Merry, B. J. (1985). The control of puberty in the dietary restricted female rat. Mechanisms of ageing and development, 32(2-3), 179–191. 10.1016/0047-6374(85)90078-8

Hubert, M. F., Laroque, P., Gillet, J. P., & Keenan, K. P. (2000). The effects of diet, ad libitum feeding, and moderate and severe dietary restriction on body weight, survival, clinical pathology parameters, and cause of death in control Sprague-Dawley rats. Toxicological Sciences, 58(1), 195–207. 10.1093/toxsci/58.1.195

Johnson, A. W. (2018). Characterizing ingestive behavior through licking microstructure: Underlying neurobiology and its use in the study of obesity in animal models. International Journal of Developmental Neuroscience, 64, 38–47. 10.1016/j.ijdevneu.2017.06.012

Kowaleski-Jones, L., Wen, M., & Fan, J. X. (2019). Unpacking the paradox: testing for mechanisms in the food insecurity and BMI association. Journal of Hunger & Environmental Nutrition, 14(5), 683–697. 10.1080/19320248.2018.1464997

Kral, T. V., Chittams, J., & Moore, R. H. (2017). Relationship between food insecurity, child weight status, and parentLreported child eating and snacking behaviors. Journal for Specialists in Pediatric Nursing, 22(2), e12177. 10.1111/jspn.12177

Leger, M., Quiedeville, A., Bouet, V., Haelewyn, B., Boulouard, M., Schumann-Bard, P., & Freret, T. (2013). Object recognition test in mice. Nature protocols, 8(12), 2531–2537. 10.1038/nprot.2013.155

Lin, W. C., Liu, C., Kosillo, P., Tai, L. H., Galarce, E., Bateup, H. S., … & Wilbrecht, L. (2022). Transient food insecurity during the juvenile-adolescent period affects adult weight, cognitive flexibility, and dopamine neurobiology. Current Biology, 32(17), 3690–3703. 10.1016/j.cub.2022.06.089 External Link

McCurdy, K., Gans, K. M., Risica, P. M., Fox, K., & Tovar, A. (2022). Food insecurity, food parenting practices, and child eating behaviors among low-income Hispanic families of young children. Appetite, 169, 105857. 10.1016/j.appet.2021.105857

Metallinos-Katsaras, E., Must, A., & Gorman, K. (2012). A longitudinal study of food insecurity on obesity in preschool children. Journal of the Academy of Nutrition and Dietetics, 112(12), 1949–1958. 10.1016/j.jand.2012.08.031

Myers, K. P., Majewski, M., Schaefer, D., & Tierney, A. (2022). Chronic experience with unpredictable food availability promotes food reward, overeating, and weight gain in a novel animal model of food insecurity. Appetite, 176, 106120. 10.1016/j.appet.2022.106120

Nettle, D., Andrews, C., & Bateson, M. (2017). Food insecurity as a driver of obesity in humans: The insurance hypothesis. Behavioral and Brain Sciences, 40, e105. 10.1017/S0140525X16000947

Nettle, D., Joly, M., Broadbent, E., Smith, C., Tittle, E., & Bateson, M. (2019). Opportunistic food consumption in relation to childhood and adult food insecurity: an exploratory correlational study. Appetite, 132, 222–229. 10.1016/j.appet.2018.07.018

Ortiz-Marrón, H., Ortiz-Pinto, M. A., Urtasun Lanza, M., Cabañas Pujadas, G., Valero Del Pino, V., Belmonte Cortés, S., … & Ordobás Gavín, M. (2022). Household food insecurity and its association with overweight and obesity in children aged 2 to 14 years. BMC Public Health, 22(1), 1930. 10.1186/s12889-022-14308-0

Power, M. L., & Schulkin, J. (2008). Anticipatory physiological regulation in feeding biology: cephalic phase responses. Appetite, 50(2-3), 194–206. 10.1016/j.appet.2007.10.006

Rasmusson, G., Lydecker, J. A., Coffino, J. A., White, M. A., & Grilo, C. M. (2019). Household food insecurity is associated with binge eating disorder and obesity. International Journal of Eating Disorders, 52(1), 28–35. 10.1002/eat.22990DigitalObject Identifier (DOI)

Robinson, M. J., Anselme, P., Fischer, A. M., & Berridge, K. C. (2014). Initial uncertainty in Pavlovian reward prediction persistently elevates incentive salience and extends sign-tracking to normally unattractive cues. Behavioural brain research, 266, 119–130. 10.1016/j.bbr.2014.03.004

Robinson, M. J., Bonmariage, Q. S. A., & Samaha, A. N. (2023). Unpredictable, intermittent access to sucrose or water promotes increased reward pursuit in rats. Behavioural Brain Research, 453, 114612. 10.1016/j.bbr.2023.114612

Rizzoto, G., Sekhar, D., Thundathil, J. C., Chelikani, P. K., & Kastelic, J. P. (2019). Calorie restriction modulates reproductive development and energy balance in pre-pubertal male rats. Nutrients, 11(9), 1993. 10.3390/nu11091993

Skvortsova, A., Veldhuijzen, D. S., Kloosterman, I. E., Pacheco-López, G., & Evers, A. W. (2021). Food anticipatory hormonal responses: A systematic review of animal and human studies. Neuroscience & Biobehavioral Reviews, 126, 447–464. 10.1016/j.neubiorev.2021.03.030

Smeets, P. A., Erkner, A., & De Graaf, C. (2010). Cephalic phase responses and appetite. Nutrition reviews, 68(11), 643–655. 10.1111/j.1753-4887.2010.00334.x

Spaulding, M. O., Hoffman, J. R., Madu, G. C., Lord, M. N., Iizuka, C. S., Myers, K. P., & Noble, E. E. (2024). Adolescent food insecurity in female rodents and susceptibility to diet-induced obesity. Physiology & Behavior, 273, 114416. 10.1016/j.physbeh.2023.114416

Stinson, E. J., Votruba, S. B., Venti, C., Perez, M., Krakoff, J., & Gluck, M. E. (2018). Food insecurity is associated with maladaptive eating behaviors and objectively measured overeating. Obesity, 26(12), 1841–1848. 10.1002/oby.22305

Tran, D., & Westbrook, R. F. (2018). Dietary effects on object recognition: The impact of high-fat high-sugar diets on recollection and familiarity-based memory. Journal of Experimental Psychology: Animal Learning and Cognition, 44(3), 217. https://psycnet.apa.org/doi/10.1037/xan0000170

Varady, K. A., Cienfuegos, S., Ezpeleta, M., & Gabel, K. (2022). Clinical application of intermittent fasting for weight loss: progress and future directions. Nature Reviews Endocrinology, 18(5), 309–321. 10.1038/s41574-022-00638-x

Wald, H. S., & Grill, H. J. (2019). Individual Differences in Behavioral Responses to Palatable Food or to Cholecystokinin Predict Subsequent DietLInduced Obesity. Obesity, 27(6), 943–949. 10.1002/oby.22459Digital Object Identifier(DOI)

Welton, S., Minty, R., O’Driscoll, T., Willms, H., Poirier, D., Madden, S., & Kelly, L. (2020). Intermittent fasting and weight loss: Systematic review. Canadian Family Physician, 66(2), 117–125. http://www.ncbi.nlm.nih.gov/pmc/articles/pmc7021351/

